# Mapping virulence-associated protein interaction networks reveals regulators of thermotolerance in *Cryptococcus neoformans*

**DOI:** 10.64898/2026.09.25.754343

**Authors:** Mayara Barbara da Silva, Mariana Cavalcante Almeida Sa, Florence Roux-Dalvai, Arnaud Droit, Jennifer Geddes-McAlister

**Affiliations:** Molecular and Cellular Biology Department, University of Guelph, Guelph, ON, Canada; Biology Department, Belhaven University, Jackson, MS, United States; Department of Molecular Medicine, Faculty of Medicine, Université Laval, Québec, QC, Canada; Proteomics and Computational Biology Laboratory, CHU de Québec - Université Laval Research Center, Québec City, Québec G1 V 4G2, Canada; Inria, Maasai team-Université Côte d’Azur, Nice, France; Canadian Artificial Intelligence and Mass Spectrometry for Systems Biology (CAN-AIMS) Consortium, Canada

**Keywords:** *Cryptococcus neoformans*, Protein-protein interactions, protein correlation profiling, heat-shock proteins

## Abstract

Protein-protein interactions (PPIs) influence critical biological processes in pathogenic microorganisms, such as the human fungal pathogen, *Cryptococcus neoformans*. Fungal thermotolerance and stress response pathways are key virulence determinants that directly impact pathogen adaptation and survival and the infection process. To establish a comprehensive baseline of PPIs in *C. neoformans* and explore these interactions to infer functional roles for uncharacterized proteins, we applied size exclusion chromatography coupled with mass spectrometry to the secreted and cellular proteomes of the fungi. As a result, 216 and 1699 unique proteins were identified across 24 secretome and proteome fractions, respectively. The predicted secretome networks included expected proteins associated with vesicles and virulence, indicating a role in extracellular defense. Whereas the cryptococcal proteome highlighted interactions among proteins with defined roles in fungal virulence for protein stability and thermotolerance, including two previously uncharacterized proteins, CNAG_00287 and CNAG_05199, putatively involved in complex formation with heat-shock proteins (HSP). Based on sequence and structure homology, we propose that CNAG_00287 is a tetratricopeptide repeat-containing co-chaperone that modulates Hsp 70 activity and CNAG_05199 functions as a Hsp70. We validated the thermotolerance role of CNAG_00287 in heat-related stress, as its absence significantly impaired fungal growth in nutrient-limited media at 37 °C. Together, this work resolves virulence-associated PPIs within *C. neoformans* and reveals new molecular regulators of thermotolerance that underpin fungal pathogenicity.

## Introduction

*Cryptococcus neoformans* is the main etiological agent of cryptococcosis, a life-threatening fungal disease with global distribution that affects approximately 200,000 people annually (1). Within immunocompromised individuals, inhaled fungal spores or desiccated cells acquired from the environment migrate from the lungs to the central nervous system (CNS), crossing the blood-brain barrier (BBB), to cause meningoencephalitis. The elaboration of cryptococcosis within immunocompromised individuals, such as HIV/AIDS patients, displays high mortality rates (up to 24%) despite therapeutic treatment (2,3). These high mortality rates are influenced by key virulence factors produced by *C. neoformans*, including the formation of a polysaccharide capsule to evade macrophage phagocytosis, production of melanin to combat reactive oxygen species, and the release of extracellular enzymes for tissue degradation and nutrient acquisition (4). Importantly, the ability to survive at human body temperature (i.e., thermotolerance), which often depends on members of the heat-shock proteins (HSP) 70 family (5,6), is a distinguishing factor of *C. neoformans*, as it enables invasion and survival within the human host.

HSPs, a well-conserved family of chaperones, drive protein folding and clearance of misfolded proteins in the cell, particularly under high-temperature stress conditions (7). In the absence of HSP pathway activation, insoluble protein aggregates accumulate within the fungal cell, causing toxicity and reducing pathogen fitness and survival (8,9). The structure of Hsp70s includes a N-terminal nucleotide-binding domain (NBD), which is responsible for the ATP binding and hydrolysis, linked to a substrate-binding domain (SBD), which binds to the specific protein’s clients. In the native form, Hsp70 domains are in a docked ADP-bound state with low substrate-affinity, and upon ATP binding, the domains undock, allowing the SBD to strongly bind to the substrate (10). Moreover, Hsp70s often use co-chaperones, such as tetratricopeptide repeat (TPR)-containing proteins and other members of the HSP family (e.g., Hsp40, Hsp70, and Hsp90), to increase ATPase activity and overall efficiency to promote pathogen stability and adaptation within the environment of the host (7).

Proteome profiling of *C. neoformans* provides critical evidence into the role of proteins regulating the production of these virulence factors, promoting fungal survival under altered environmental conditions, and modulating antifungal resistance (11–14). Additionally, investigation into *Cryptococcus* spp. protein-protein interactions (PPIs) or protein complexes provide biological insight into patterns of cellular regulation, signaling cascades, and downstream activation processes (15,16). Critically, many strategies for detecting and characterizing PPIs require *a priori* knowledge of the interactors (i.e., bait), limiting unbiased detection of PPIs within a single or multiple biological systems. For instance, identification of a single protein of interest and the addition of an exogenous tag to the protein, may disrupt biologically relevant interactions (15). Moreover, assigning a specific bait protein substantially restricts the breadth of investigations to a single candidate and its interactors, potentially missing other interactions driving processes within a cell (16,17). Importantly, recent applications of co-fractionation methods, such as size exclusion chromatography coupled to mass spectrometry (SEC-MS), have characterized proteoforms and protein complexes in an unbiased manner (18–20), including defining signature responses and inhibition of HSPs (21). These approaches provide a key advantage of exerting minimal impact on interacting systems as the porous column used for SEC separates analytes based on their molecular weight (16,22) and outputs protein correlation profiles for interpretation.

In this study, we applied SEC-MS to resolve virulence-associated PPIs across the secreted and cellular proteomes of *C. neoformans*. We identified 216 and 1,699 unique proteins across fractionated secretome and cellular datasets, respectively. The secretome analysis revealed extracellular networks enriched in vesicle-associated virulence factors, whereas the cellular proteome defined intracellular interactomes linked to protein stability and thermotolerance. Notably, we identified two previously uncharacterized proteins, CNAG_00287 and CNAG_05199, predicted to form complexes and stabilize HSPs. Structural and sequence analyses suggest CNAG_00287 functions as a tetratricopeptide repeat-containing co-chaperone modulating Hsp70 activity, while CNAG_05199 acts as an Hsp70-like protein. Within this study, we focused on CNAG_00287 for experimental validation with functional assays confirming that loss of CNAG_00287 reduces fungal growth under heat stress. These findings establish a role for CNAG_00287 in thermotolerance and highlight the utility of PPI mapping for uncovering regulators of complex biological systems. Together, these findings establish a secretome and cellular proteome map of virulence-associated interactomes that we leverage to uncover molecular regulators of fungal pathogenicity.

## Materials and Methods

### Cell culture, lysis, and protein extraction

*Cryptococcus neoformans var. grubii* wild-type (WT) strain H99 (serotype A) was maintained on yeast peptone dextrose (YPD) agar plates (2% dextrose, 2% peptone, 1% yeast extract, 1% agar) at 30 °C. For co-fractionation assays, *C. neoformans* H99 was cultured overnight at 30 °C in YPD and sub-cultured at 1:5 in yeast nitrogen base (YNB) for approximately 16 h at 37 °C. For secretome samples, supernatant of culture was collected and filtered with 0.2-µm syringe filter to remove cell debris (23). Cellular samples were collected and washed twice in phosphate buffered saline (PBS), then resuspended in M-PER™ Mammalian Protein Extraction Reagent (Thermo Scientific) containing the cOmplete™ Protease Inhibitor Cocktail (Roche) followed by probe sonication in an ice bath for 5 cycles of 30 s on and 30 s off (power 30%) (23). After lysis, the samples were centrifuged at 10,000 RCF for 10 min to precipitate cell debris. Protein quantification was performed using Pierce™ BCA Protein Assay Kits (Thermo Scientific) for both secretome and proteome analysis to ensure uniform protein loading during co-fractionation assays.

### Protein co-fractionation by size exclusion chromatography

Protein fractionation was performed using the NGC fast protein liquid chromatography system (Bio-Rad) with a Superose 6 Increase 10/300 SEC column (Cytiva). Protein extracts (approximately 80-100 µg of total protein) were resolved into 28 fractions consisting of 1 mL, of which 24 were selected based on A206/A280 chromatogram peaks. Overnight digestion of selected fractions was performed using LysC/trypsin (25:1 protein:enzyme ratio), then stopped by the addition of a 20% acetonitrile/6% trifluoroacetic acid solution.

### Mass spectrometry instrument operation

The digested samples were purified using the EvoSep One high-performance liquid chromatography (Evosep Biosystems) system and analyzed on an Orbitrap™ Exploris 480 (Thermo Scientific) mass spectrometer. The chromatographic separation was performed using a manufacturer-defined 100 samples per day method allowing peptide elution with a 11.5 min gradient of 0.1% formic acid in acetonitrile at a flow rate of 1 μL/min on an Evosep EV1106 separation column (8 cm long, 150 μm internal diameter and 1.5 μm beads). The mass spectrometer operated in the data-independent acquisition mode, in which full scan mass spectra (300 to 1500 *m*/*z*) were acquired at a resolution of 60,000 with a normalized automatic gain control (AGC) of 300% and a maximum injection of 118 ms. Each mass spectrometry scan included 18 *m/z* windows across the 350-872 *m*/*z* mass range with fragmentation by Higher energy Collision induced Dissociation (HCD) at 30% of normalized collision energy. The resulting fragments were detected in the Orbitrap at a resolution of 15,000, with a normalized AGC of 800% and a maximum injection time of 22 ms.

### Mass spectrometry data processing

Mass spectrometer raw files were searched using DIA-NN for the secretome and Spectronaut v.19 (Biognosys) for the proteome, with protein FASTA files against *C. neoformans* var. *grubii* serotype A (strain H99; UP000010091; 7,429 sequences) retrieved from UniProt (https://www.uniprot.org). Set parameters included trypsin enzyme specificity at a maximum of two missed cleavages, carbamidomethylation of cysteine as fixed modification, and methionine oxidation and N-acetylation as variable modifications. False Discovery Rate (FDR) was set to 1% and a minimum of two peptides per protein were required for protein identification.

Processed files were analyzed and visualized in Perseus (version 1.6.2.2) (24) filtered for contaminants and duplicates. Proteins intensities were log_2_ transformed, normalized, and valid values were filtered for presence in at least 50% of replicates for secretome and at least 70% of replicates for cellular proteome. The size distribution for each fraction was determined via the averaged molecular mass proteins within the fraction.

### Protein interaction network prediction

Protein complexes were predicted using the STRING network database (www.string-db.org; STRG0A93OCR uploaded proteome obtained from UniProt) for each well-resolved fraction. Unresolved fractions (< 5 kDa average size difference) were combined prior to prediction to minimize false negatives, and proteins consistently identified in all fractions were removed from subsequent analysis. Complexes were curated for physical subnetwork, medium confidence interaction score (0.400), and 1% FDR. Clustering was defined applying the Markov Cluster (MCL) Algorithm (inflation parameter: 3) (25). Networks were reconstructed and visualized in Cytoscape 3.10.4 (26).

### Sequence and structure analyses

Prioritized interacting structures were modelled using AlphaFold 3.0 and ranked according to Interface Predicted Template Modelling (ipTM), inter-chain Predicted Aligned Distance (PAE), and Predicted Local Distance Difference Test (pLDDT) (27). Structures presenting an ipTM > 0.4, PAE < 5 Å, and pLDDT > 70 were considered of high confidence and were forwarded to further analyses (27). Multiple sequence alignments were performed using ClustalW and visualized in Espript 3.2 (28). Maximum-likelihood phylogenetic trees were created using PhyML 3.1, with sequence alignment obtained via MUSCLE 3.8.31 and refined by Gblocks 0.91b (29). Additionally, superimpositions to crystal structures obtained from the Protein Data Bank (PDB) were performed using the DALI server (30). All complexes were visualized using ChimeraX 1.11.1 (31).

### Thermotolerance assay

*Cryptococcus neoformans* KN99 WT and CNAG_00287Δ strains (32), were cultured overnight in YPD at 30 °C, followed by a 1:100 subculture in YPD or YNB for approximately 16 h at 30 °C. Cells were diluted to a OD_600nm_ of 0.1 and incubated in 96-well plates at 30 and 37 °C for 48 h in both media. The experiment was performed in five biological replicates, with OD_600nm_ recorded every 15 min. GraphPad Pris v.10.6.0 was used to visualize the growth curve graph, calculate the area under the curve, and perform unpaired two-way t-tests to determine statistical significance (p < 0.05).

## Results

### SEC fractionation coupled with mass spectrometry reveals a global perspective of the interactome

*Cryptococcus neoformans* secretome (i.e., supernatant) and proteome (i.e., cell lysate) fractionation yielded 24 co-elution fractions per replicate. For the secretome, elution profiles were as expected, with the size of proteins within the SEC fractions decreasing across the elution gradient **(Figure 1A).** A total of 216 unique proteins were identified across six fractions, with the highest number of proteins in fractions 16 and 17, accounting for 157 and 181 identifications, respectively **(Figure 1B)**. These fractions shared 122 proteins, with 30% and 20% of identifications being unique to each, respectively **(Figure 1C)**.

**Figure 1.**
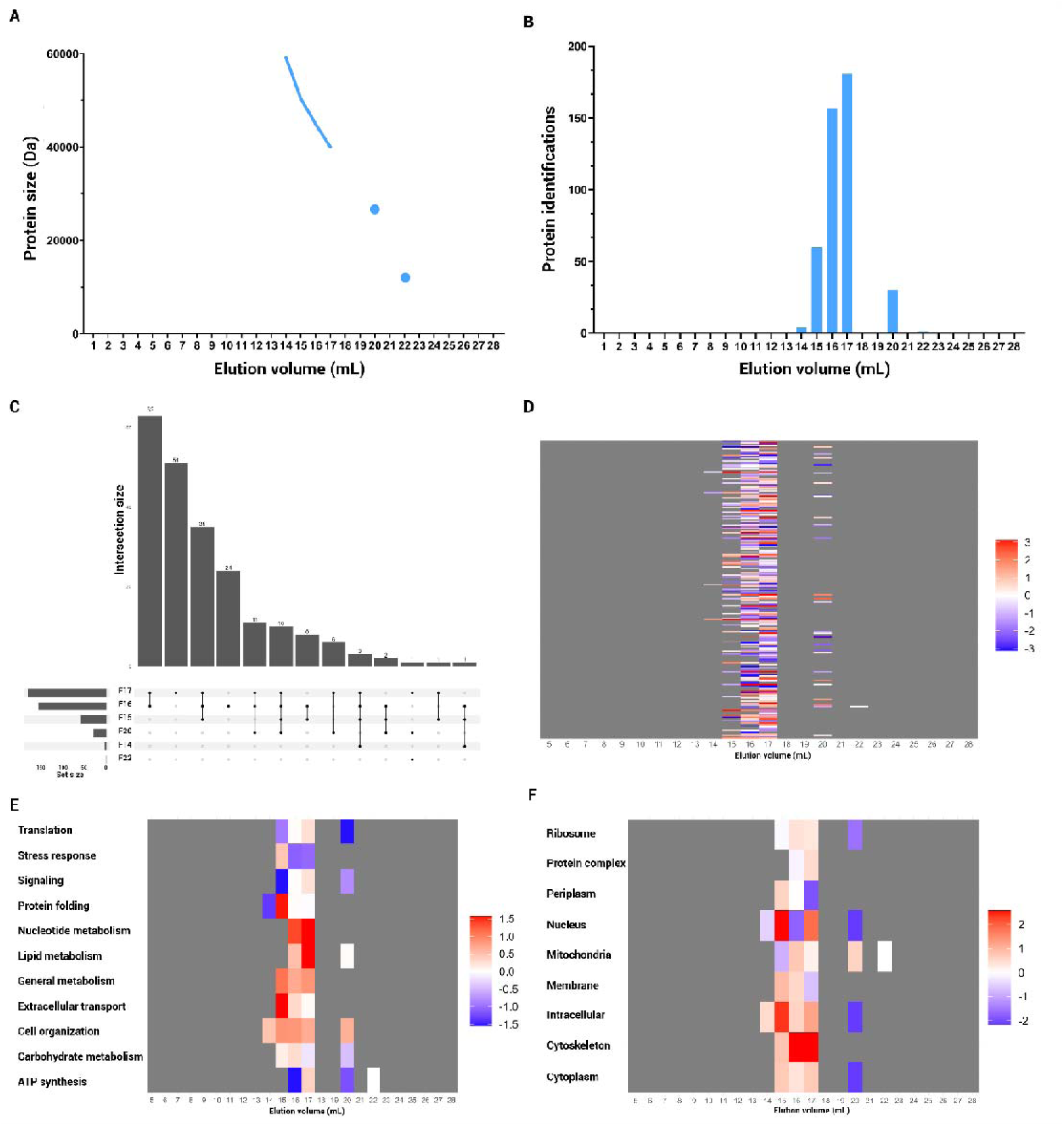
SEC protein abundance profiles of *C. neoformans* secretome. **A,** Fractionation profile highlighting protein size distribution per collected fraction (1 mL). Fraction size was determined by averaging the molecular mass of protein identified in each fraction. **B,** Number of proteins identified per fraction. **C,** Upset plot representing the number of proteins shared across specific fractions. **D,** Protein intensities distributed per fraction. **E,** Gene Ontology Biological Processes (GOBP) analyses depicting key enriched functions. **F,** Gene Ontology Cellular Compartment (GOCC) analyses showing cellular localization enrichment. Heat-map intensities represent the average of median-normalized values. Experiment performed in biological quadruplicate.

Normalized intensity values of secretome proteins were mostly consistent across fractions, with fractions 16 and 17 displaying the highest abundances **(Figure 1D)**. Notably, highly abundant proteins included the HSPs, SSA1 and thioredoxin. In addition, proteins associated with metabolic functions (e.g., lipid and nucleotide metabolism) were consistently more abundant across fractions, whereas proteins associated with extracellular transport, protein folding, and stress response were mostly present in fraction 15 **(Figure 1E)**. Secretome fractions also contained proteins from diverse cellular localizations, including intracellular organelles, the cytoskeleton, and the cytoplasm **(Figure 1F)**.

For cellular proteome fractionation, we also observed the anticipated size pattern across fractions 6-22, but with notable fluctuation in the early and late fractions **(Figure 2A)**. These fluctuating size values likely correspond to low detectable protein levels (approx. 10% of total identification) within the tail fractions **(Figure 2B)**. Overall, 1698 unique proteins were identified, with approximately 1000 proteins detected in fractions 6-18, accounting for the bulk of the cellular proteome **(Figure 2B)**. Approximately 35% of proteins identified in the first half of the elution process (i.e., fractions 6 to 18) were consistently found across these fractions. This pattern was likely due to the high protein abundance in the cell lysate, which led to a large number of identifications in these fractions **(Figure 2C and 2D, Figure S1)**.

**Figure 2.**
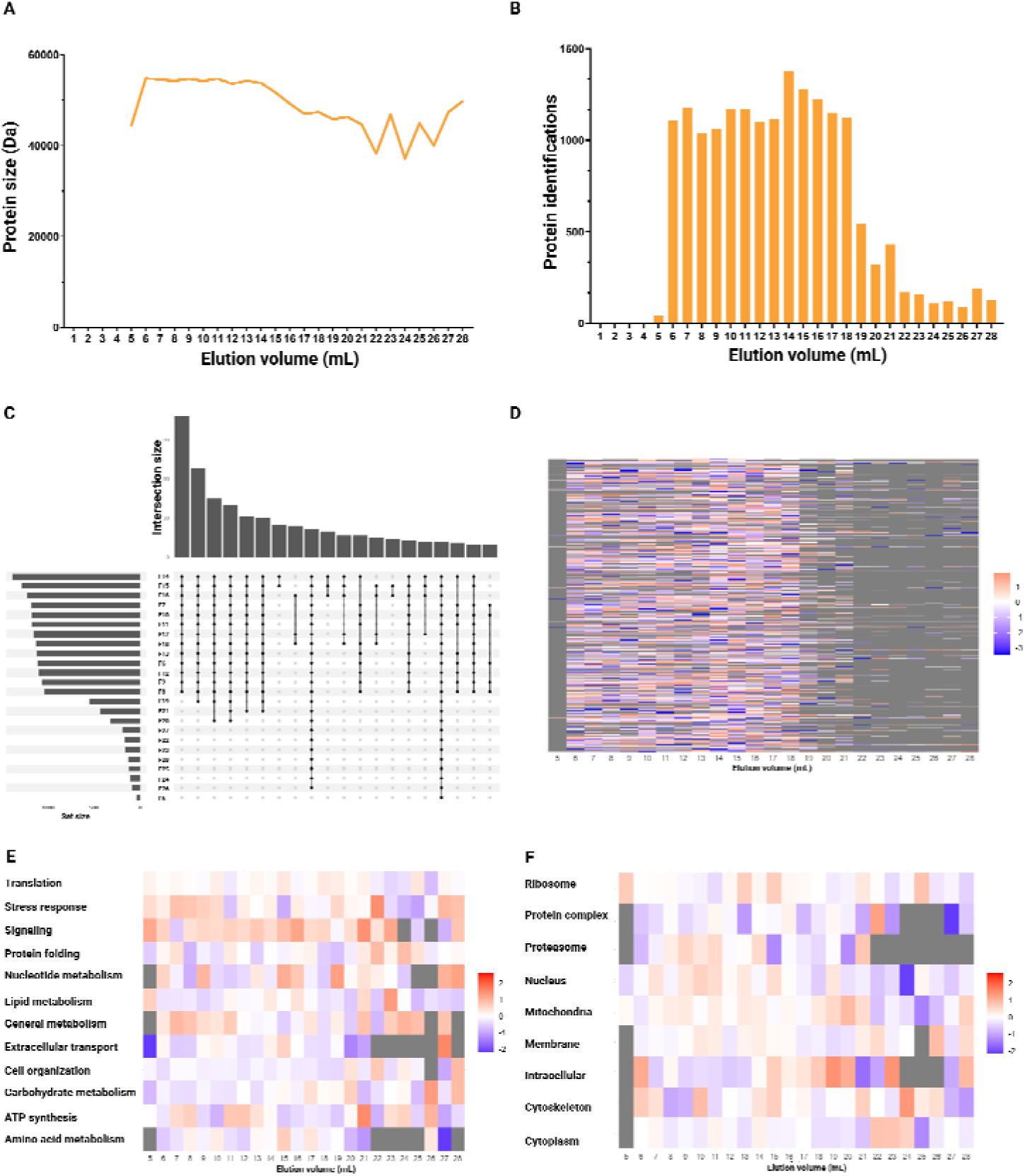
SEC protein abundance profiles of *C. neoformans* cellular proteome. **A,** Fractionation profile highlighting protein size distribution per collected fraction (1 mL). Fraction size was determined by averaging the molecular mass of the detected proteins. **B,** Number of proteins identified per fraction. **C,** Upset plot representing the number of proteins shared across specific fractions. **D,** Protein intensities distributed per fraction. **E,** GOBP analysis depicting key enriched functions. **F,** GOCC analysis showing cellular localization enrichment. Heat-map intensities represent the average of median-normalized values. Experiment performed in biological quadruplicate.

We observed that cryptococcal proteome proteins detected in multiple fractions displayed high variability in their intensity profiles. Nevertheless, higher-intensity and - frequency identification patterns were still observed in fractions 15-18, comparable to the secretome **(Figure 2C and D)**. Among the most abundant proteins in the cellular proteome were HSP 9/12 and CipC. Moreover, stress response and general metabolic functions were enriched in the initial and final fractions, while signalling pathways were highly abundant across all fractions **(Figure 2E)**. The cellular proteome also contained proteins localized to several compartments (e.g., ribosomes and mitochondria); however, no clear enrichment distribution was observed regarding cellular localization and eluted fractions **(Figure 2F)**.

### Curated protein networks highlight key clusters and biological functions in the secretome

After establishing fractionation patterns of the *C. neoformans* secretome and cellular proteome, we aimed to predict co-eluted proteins that form physical complexes. Therefore, we applied STRING PPI network enrichment analysis to fractions 14-17, 20, and 22 of the cryptococcal secretome (fractions selected on protein identifications and abundance). Of these six fractions, only 16 and 17 showed PPI enrichment; the remaining fractions lacked sufficient identifications to generate reliable networks. Overall, proteins clustered by biological function, highlighting the correlation between co-localization and biological processes. Specifically, in fraction 16, eight functionally annotated clusters associated with ATP generation, protein translation, and folding were identified, with the latter demonstrating the highest enrichment signal **(Figure 3A and 3B).** Given the role of thermotolerance in cryptococcal virulence, we focused on identification of HSPs and their affiliated proteins within the interaction networks. Notably, the chaperones SSA1, Hsp70, and Hsp90 were predicted to be key in the protein folding cluster, forming several interactions. In the subsequent fraction 17, nine clusters were identified. In addition to the functions reported in fraction 16, fraction 17 displayed nucleus-associated clusters and enrichment of ribonucleoside triphosphate metabolism and biosynthetic processes **(Figure 3C and D)**. Interestingly, the observed enrichment for proteins associated with protein folding was maintained across fractions despite a decrease in protein count.

**Figure 3.**
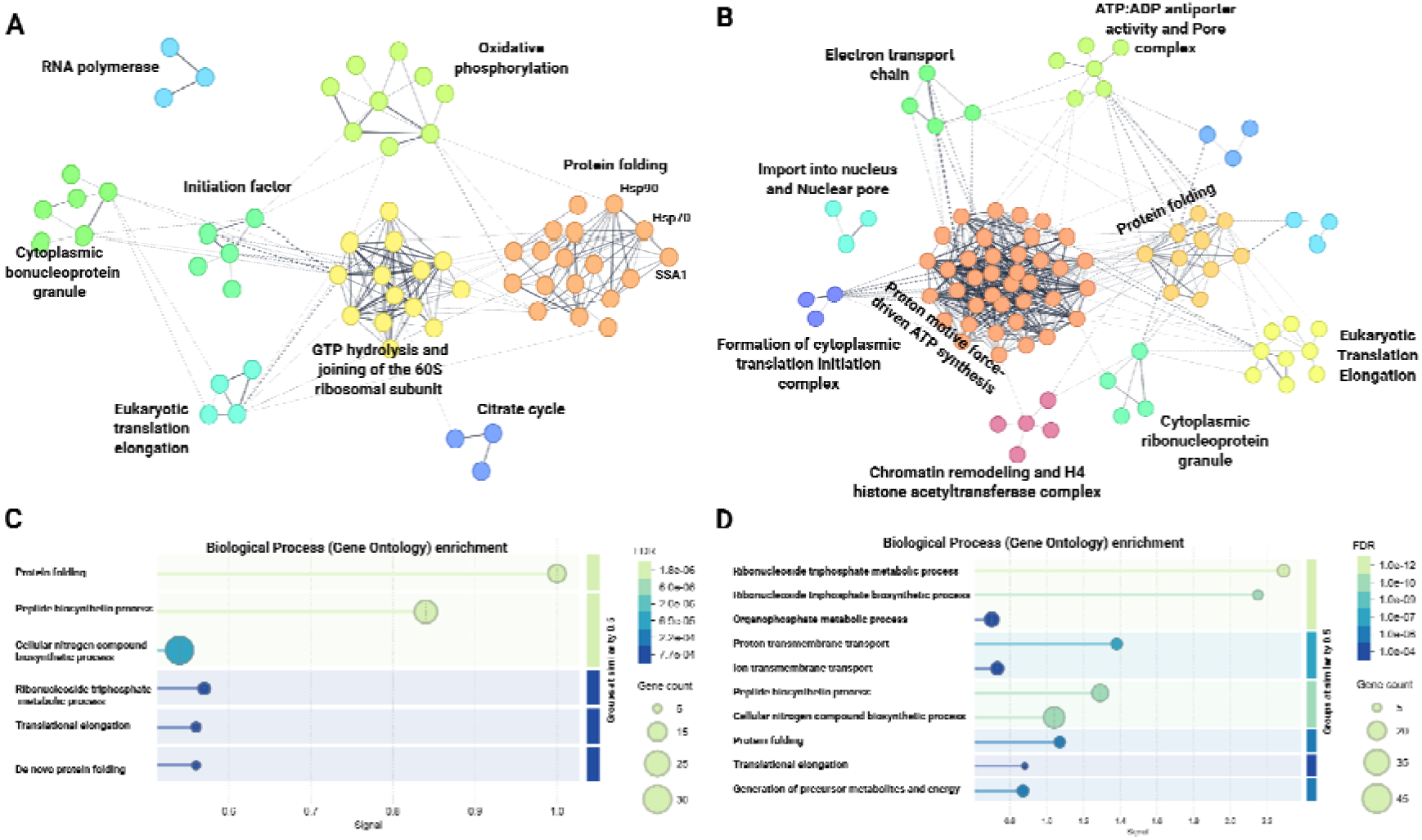
STRING interaction complex subnetwork predictions of *C. neoformans* secretome inferred from SEC-MS fractionation. **A,** Secretome PPI network obtained from fraction 16 highlighting three HSPs in the protein folding cluster. **B,** Secretome PPI network obtained from fraction 17 highlighting key metabolic clusters. **C,** GOBP enrichment of fraction 16. **D,** GOBP enrichment of fraction 17. STRING parameters: confidence interaction score = 0.400, FDR = 1%, MCL inflation parameter = 3, PPI enrichment p-value < 1.0^-16^.

### Cellular proteome interaction network prediction highlights uncharacterized proteins with putative roles in heat shock response

Building on the secretome analysis, cryptococcal cellular proteome prediction networks were generated by removing proteins consistently identified across all fractions and merging similar-sized fractions. This approach yielded five combined fractions for input into the STRING network, of which four showed PPI enrichment. Combined fractions two and three, encompassing fractions 6-15 and 16-21, respectively, exhibited comparable prediction networks. Specifically, approximately 200 clusters were identified across each fraction, with the top 20 MCL clusters and their associated enrichments being depicted (**Figure 4A).** These clusters were associated with diverse biological processes, with a high prevalence of ribosomal and mitochondrial proteins identified. Of note, ribosomal and mitochondrial-associated clusters were highly interconnected and contained nine uncharacterized proteins. General metabolic biological processes, such as organonitrogen biosynthesis and ATP generation, were highly enriched in the predicted networks **(Figure 4B).**

**Figure 4.**
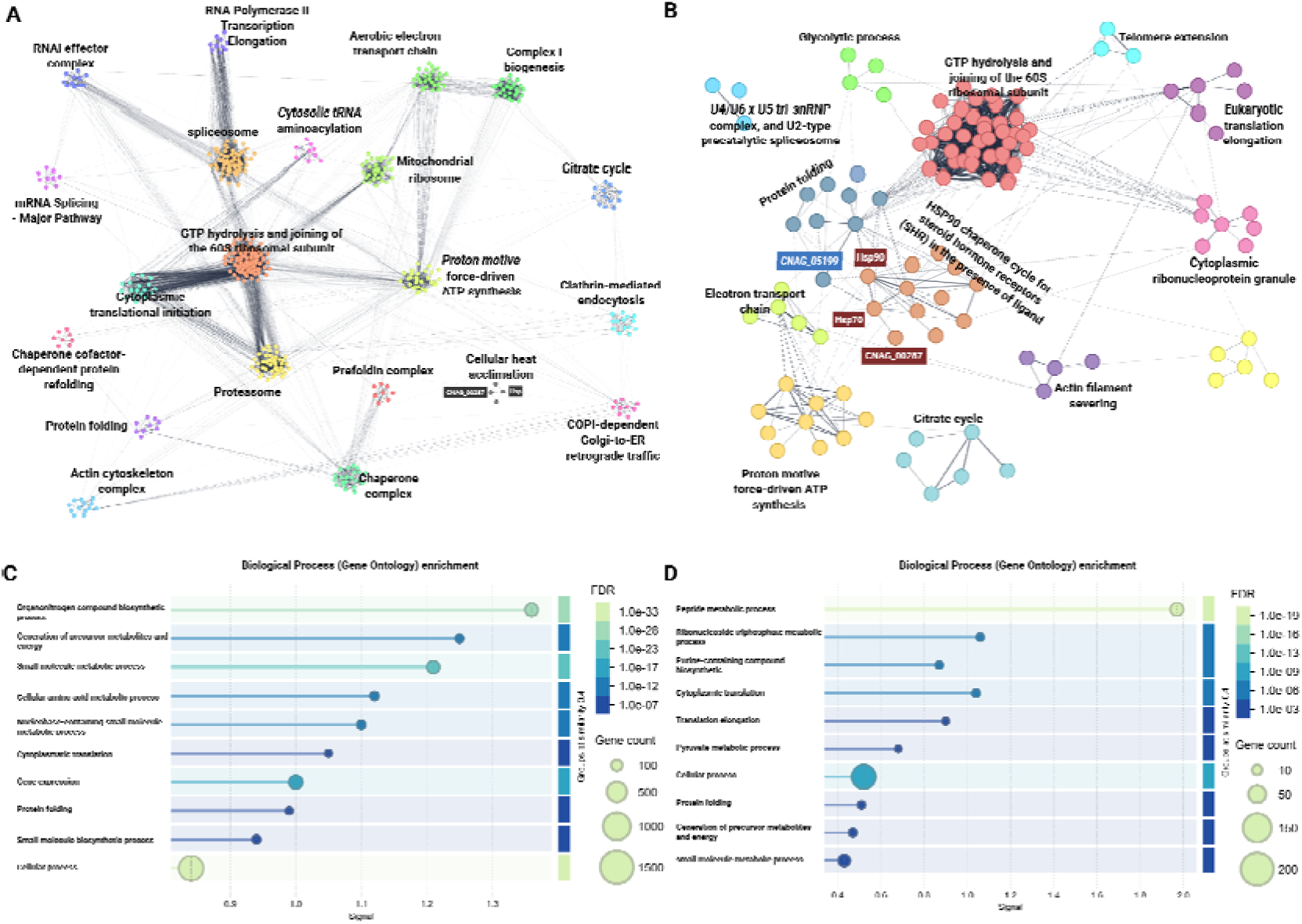
STRING interaction complex subnetwork predictions of *C. neoformans* cellular proteome inferred from SEC-MS fractionation. A, Proteome PPI network obtained from combined fraction two (fractions 6-15) highlighting top 20 clusters. **B,** Proteome PPI network obtained from combined fraction five (fractions 25-28) displaying 12 clusters. Hsp70/Hsp90 and their interacting uncharacterized proteins are highlighted. **C,** GOBP enrichment of combined fraction two. **D,** GOBP enrichment of combined fraction five. STRING parameters: confidence interaction score = 0.400, FDR = 1%, MCL inflation parameter = 3, PPI enrichment p-value < 1.0^-16^. Clusters with gene counts < 3 were not considered during analysis.

Similar to combined fractions two and three, the subsequent fractions also showed similar network profiles. For instance, combined fractions four and five, encompassing fractions 22-24 and 25-28, respectively, generated approximately 13 PPI clusters each **(Figure 4C).** Again, we focused on detection of HSP within the cellular interactome and observed that combined fraction five also highlighted the strong interplay between protein folding and the Hsp90 chaperone clusters. In addition to the known interactions among HSPs, these clusters revealed two uncharacterized proteins, CNAG_00287 and CNAG_05199. Based on the STRING PPI, CNAG_00287 was predicted to interact with Hsp70 while CNAG_05199 bridged these two clusters via interactions with Hsp90 and Hsp70. Moreover, CNAG_00287 was also predicted to interact with the cellular heat acclimation cluster and to interact with another HSP in combined fraction 2 **(Figure 4A)**. Across the interaction networks, peptide metabolic processes as defined by GOBP was the most highly enriched category (**Figure 4D**). Given the importance of HSPs in cryptococcal survival and virulence, we decided to focus further experiments on the roles of the uncharacterized proteins, CNAG_00287 and CNAG_05199, using sequence and structural homology analyses.

### Classification of CNAG_00287 as a glutamine-rich tetratricopeptide repeat-containing cytoplasmic protein by multiple sequence alignments and binding pocket structure

Based on sequence homology searching, CNAG_00287 is a putative small glutamine-rich tetratricopeptide repeat-containing (SGT) cytoplasmic protein. The multiple sequence alignment (MSA) demonstrates conservation of the tetratricopeptide repeat (TPR) domain of CNAG_00287 with SGT proteins across several biological systems, including other yeasts, filamentous fungi, and mammals (**Figure 5A**). The conserved residues and sequence features within the TPR domain support the assignment of CNAG_00287 as an SGT protein **(Figure 5A; Figure S2)**. The phylogenetic analysis further illustrates the evolutionary relationships among the SGT proteins, with CNAG_00287 clustering most closely with Sgt2, a well-characterized SGT from *Saccharomyces cerevisiae* **(Figure 5B)**. The conservation observed at the sequence level is also reflected in the predicted structure of CNAG_00287. Structural mapping of sequence conservation onto the TPR domain reveals a highly conserved region forming the binding pocket associated with recognition of the Hsp70 C-terminal tail **(Figure 5C)**.

**Figure 5.**
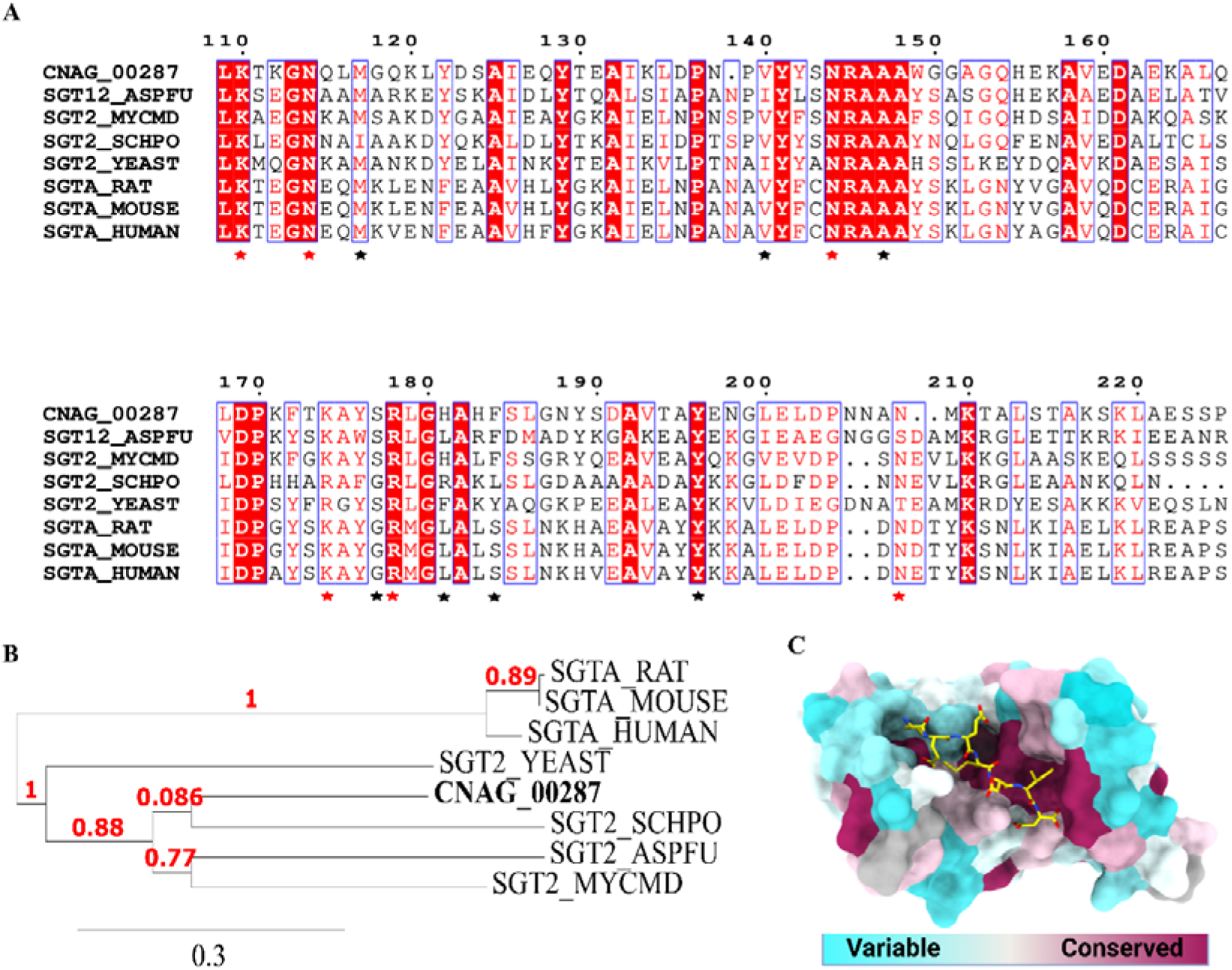
Homology analyses of CNAG_00287 and small glutamine-rich tetratricopeptide repeat-containing (SGT) proteins. **A,** Multiple sequence alignment analysis depicting high conservation (red residues and background) in the TPR domain (stars). Red stars represent the carboxylate clamp, and black stars represent the residues responsible for stabilizing the interaction. **B,** Phylogenetic tree highlighting high similarities among SGTs. **C,** Structure representation of CNAG_00287 TPR domain sequence conservation highlighting the conserved binding pocket. The Hsp70 C-terminal tail ligand is depicted in yellow. SGTA_RAT: *Rattus norvegicus;* SGTA_MOUSE: *Mus musculus;* SGTA_HUMAN: *Homo sapiens;* SGT2_YEAST: *Saccharomyces cerevisiae;* SGT2_SCHPO: *Schizosaccharomyces pombe;* SGT12_ASPFU: *Aspergillus fumigatus;* SGT2_MYCMD: *Mycosarcoma maydis*.

### Prediction of interaction between CNAG_00287 and Hsp70 supports classification as an HSP co-chaperone

In *C. neoformans,* structure prediction of the CNAG_00287-Hsp70 complex displayed a moderately confident interaction (ipTM = 0.32, inter-chain PAE; 5.38-6.37 Å), which we deemed to be associated with the low prediction confidence of the CNAG_00287 chain (pTM = 0.39). Despite this intermediate score, the model was confident in the CC-TPR/Tail assembly (pLDDT > 70) **(Figure 6A and 6B)**. As a result, isolating the CNAG_00287 TPR domain associated with the Hsp70 C-terminal increased the prediction to a very high confidence score (ipTM = 0.82, inter-chain PAE = 1.02-1.49 Å, pLDDT > 90) (**Figure 6C and 6D)**. This prediction also highlighted key residues in the binding cavity forming a carboxylate clamp-TPR (CC-TPR) structure that functions to trap terminal acidic residues (**Figure 6C and D)**. Moreover, superimposition of the CNAG_00287 and Sgt2 TPR domains demonstrated high structural homology, with 43% sequence identity and an RMSD of 0.9 Å over 120 Cα atoms **(Figure 6E)**. Altogether, these results corroborate the CNAG_00287-Hsp70 complex formation initially inferred from the SEC-MS profiling analysis.

**Figure 6.**
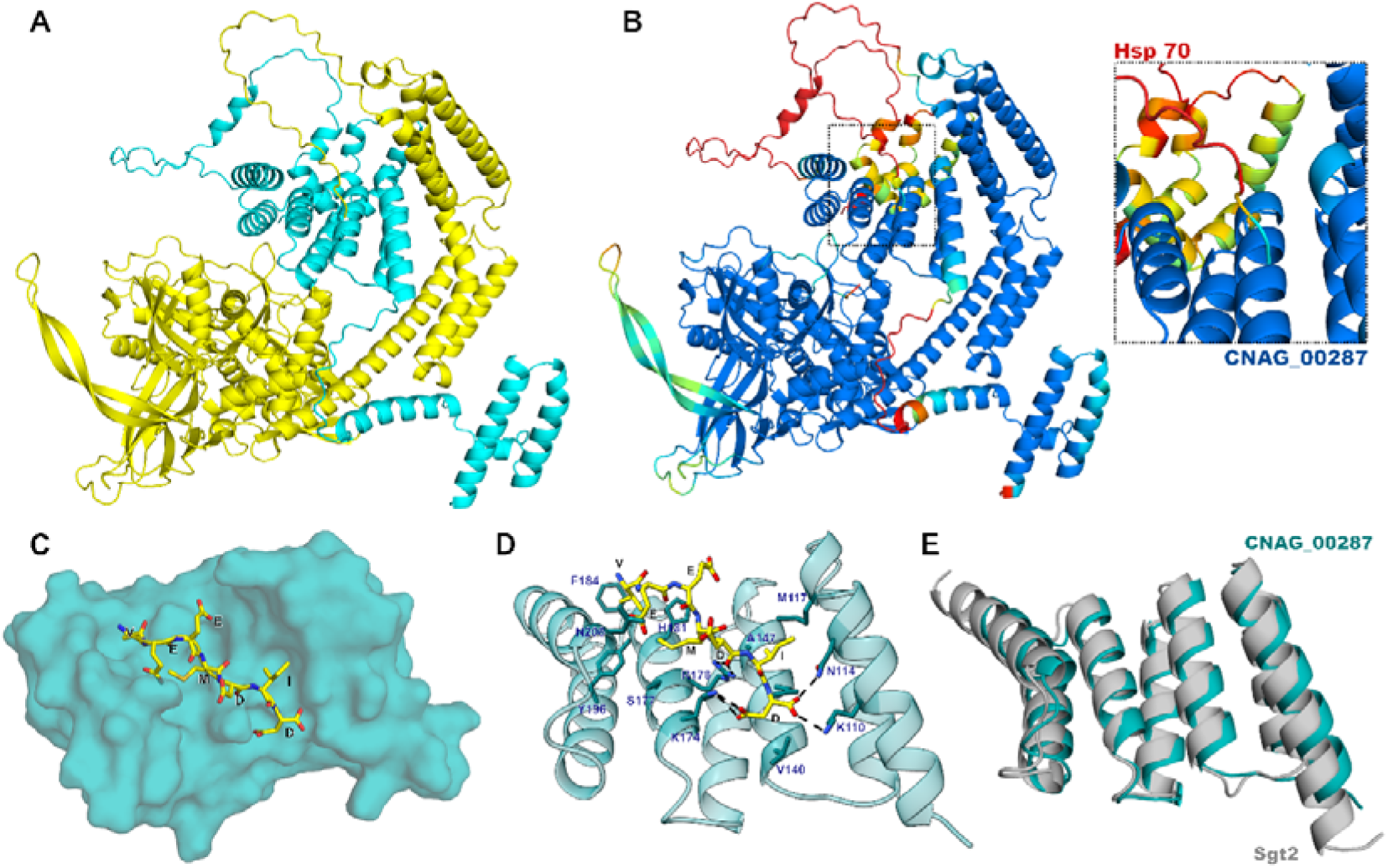
AlphaFold 3.0 structure representations of the CNAG_00287-Hsp70 predicted complex. **A,** Interacting complex colored by chain: CNAG_00287 depicted in cyan and Hsp70 depicted in yellow. **B,** Interacting complex colored by pLDDT. Confidently predicted residues are displayed in cyan (pLDDT > 70) and dark blue (pLDDT > 90), while non-confidently predicted are displayed in yellow (50 < pLDDT < 70) and red (pLDDT < 50). Magnified inset highlights the high confidence in the CC-TPR/C-terminal tail region. **C,** CNAG_00287 TPR domain (cyan) surface representation of binding cavity accommodating HSP70 C-terminal tail (yellow). **D,** Binding site highlighting key residues in CNAG_00287 (cyan; blue label) and HSP70-4 (yellow; black label). Hydrogen bonds are depicted as black dashes. **E,** Superimposition of TPR domains from predicted CNAG_00287 and Sgt2 x-ray (*Saccharomyces cerevisiae*; PDB: 5LYN) structures demonstrating high structure homology. Z-score: 21.7, RMSD 0.9 Å over 120 out of 133 Cα.

### CNAG_00287 influences thermotolerance under nutrient-limited conditions

Given its interaction with Hsp70, we predicted that CNAG_00287 may play a role in thermotolerance of *C. neoformans*. Interestingly, *C. neoformans* KN99 CNAG_00287Δ strains did not show impaired growth when subjected to a temperature increase (30 °C to 37 °C) in rich media (YPD). However, we did observe a significant reduction in growth (p < 0.0001) for the mutant strain compared to the *C. neoformans* KN99 WT strain independent of temperature, suggesting an impact of CNAG_00287 on fungal fitness **(Figure 7A and 7B)**. Next, given the important of HSPs in thermotolerance and the connection to host survival, which represents a resource-limited setting, we explored the impact of CNAG_00287Δ on thermotolerance and fungal growth under a nutrient-limited environment (i.e., YNB). Under these nutrient-limited conditions, we did not observe a significant reduction in fungal growth between the WT and CNAG_00287Δ strains at 30 °C (**Figure 7C**). However, we did observe a significant reduction in growth (supporting a defect in thermotolerance) for the CNAG_00287Δ strain compared to WT at 30 °C under the nutrient-limited conditions (**Figure 7D**). These findings support a role of CNAG_00287 in heat shock response and thermotolerance for *C. neoformans* under nutrient-limited conditions.

**Figure 7.**
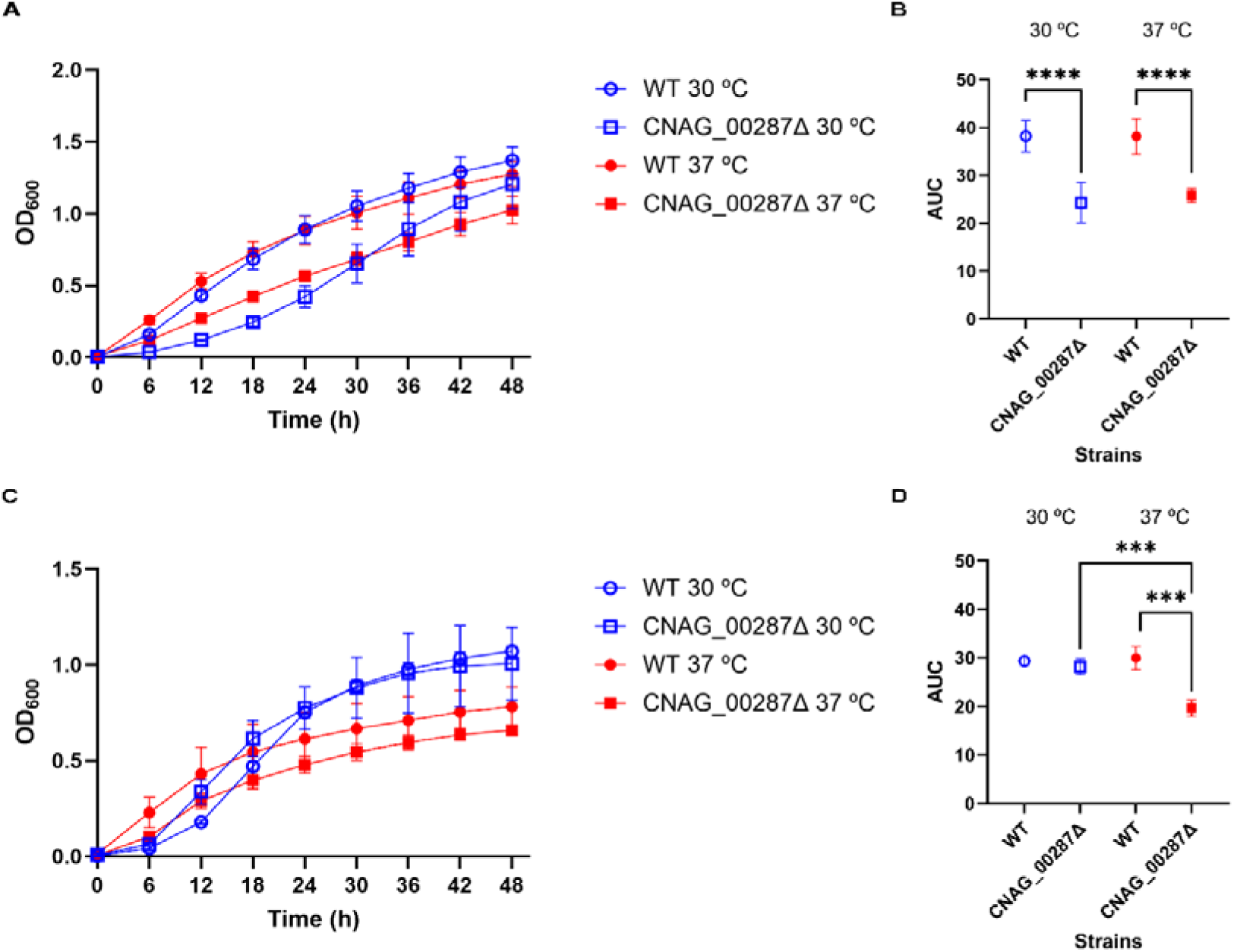
*C. neoformans* CNAG_00287Δ growth profiles in nutrient-rich (YPD) and -limited (YNB) media. **A,** Growth curve performed in YPD at 30 °C and 37 °C for 48 h. **B,** Area under the curve calculations for growth in YPD comparing WT and CNAG_00287Δ strains. Unpaired t-tests (****p < 0.0001). **C,** Growth curve performed in YNB at 30 °C and 37 °C for 48 h. **D,** Area under the curve calculations for growth in YNB comparing WT and CNAG_00287Δ strains. Unpaired t-tests (***p < 0.001) at 37 °C for CNAG_00287Δ in comparison with WT at the same temperature and CNAG_00287Δ at 30 °C. Non-significant results (p > 0.05) are not displayed. Experiment performed in biological triplicate and technical duplicate.

### CNAG_05199 is predicted to function as an Hsp70-like protein

As the predicted CNAG_00287-Hsp70 complex from SEC-MS profiling revealed a role for CNAG_00287 in stress response, we next investigated a second uncharacterized protein detected within the cellular proteome, CNAG_05199. A multisequence structure alignment of CNAG_05199 and Hsp70s revealed strong conservation of the NBD and SBD regions **(Figure 8A, Figure S3)**. Similar to *S. cerevisiae* SSQ1 (HSP7Q_YEAST), involved in mitochondrial iron-sulfur cluster biogenesis, CNAG_05199 has a short N-terminal peptide preceding the NBD. Superimposition of the predicted CNAG_05199 structure and the *Escherichia coli* Hsp70 structure, the closest homolog with an experimentally resolved crystal structure (DNAK), supported the homology inferred from sequence analysis **(Figure 8B)**. The NBD showed a Z-score of 53.2, RMSD of 1.8 Å over 383 residues and 51% identity; the SBD showed a Z-score of 25.5, RMSD of 1.9 Å over 218 residues and 47% identity. Moreover, the clustering of CNAG_05199 within the Hsp70 family, as demonstrated by phylogenetic analysis, supports its close evolutionary relationship with Hsp70 proteins from diverse organisms, including yeasts, bacteria, and mammals, as well as with a confirmed *C. neoformans* Hsp70 **(Figure 8C)**. Based on these analyses, we propose that CNAG_05199 acts as a mitochondrial Hsp70 in *C. neoformans*.

**Figure 8.**
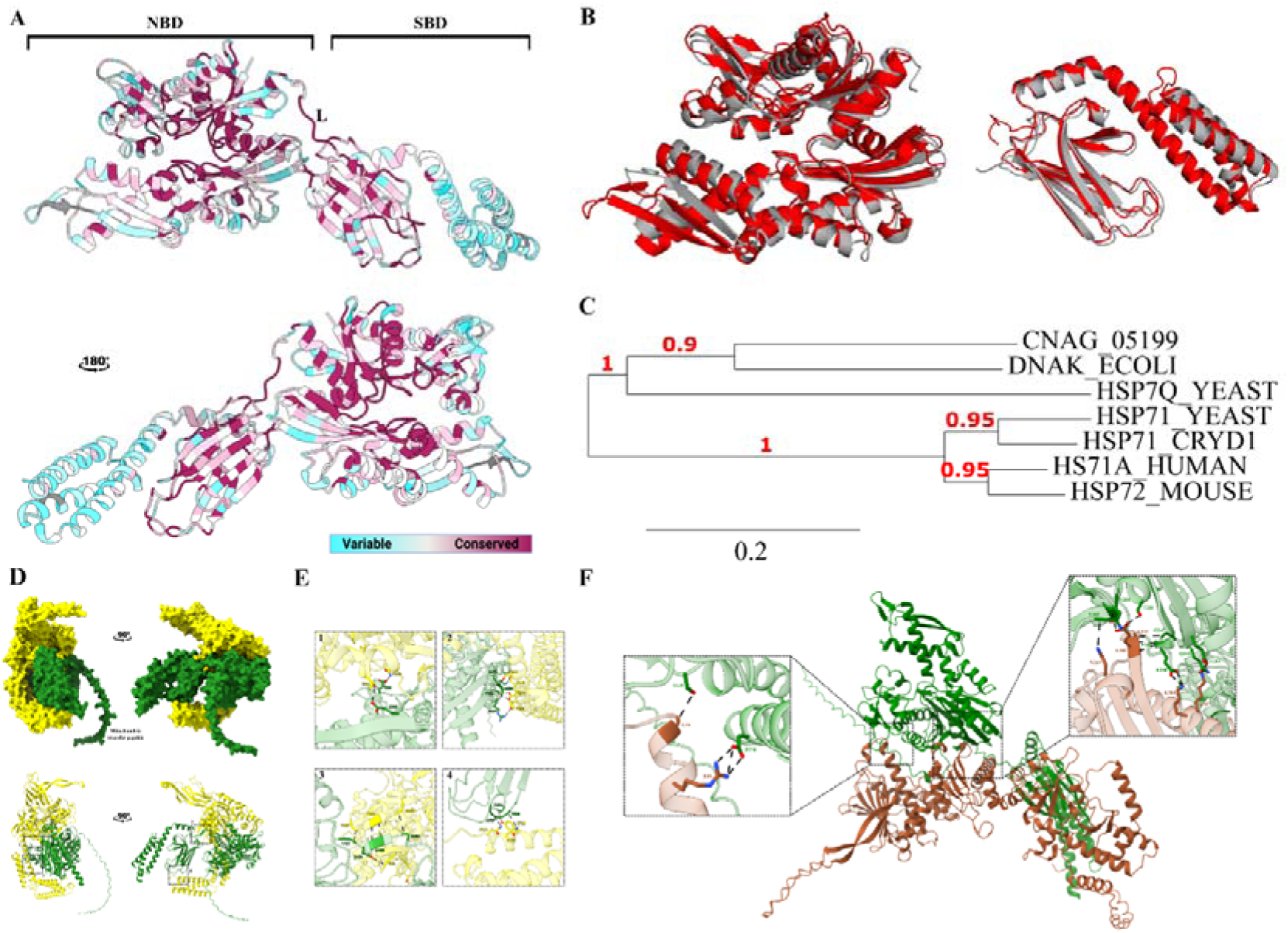
CNAG_05199 homology and structure analyses. **A,** Structural representation of ADP-bound CNAG_05199 highlighting the high degree of conservation through the whole protein. NBD: Nucleotide-binding domain; L: Interdomain linker; SBD: Substrate-binding domain. **B,** Superimposition of CNAG_05199 NBD (left) and SBD (right) to the DNAK crystal structure (PDB: 2KHO) showing high structure homology. RMSD: 1.8-1.9 Å. **C,** Maximum-likelihood phylogenetic tree demonstrating close relationship between CNAG_05199 and the Hsp70 family. DNAK_ECOLI: *Escherichia coli;* HSP7Q_YEAST and HSP71_YEAST: *Saccharomices cerevisiae,* mitochondrial and cytoplasmatic; HSP71_CRYD1: *Cryptococcus neoformans,* cytoplasmatic; HS71A_HUMAN: *Homo sapiens*; HSP72_MOUSE: *Mus musculus.* **D,** Overview of Hsp70 (yellow) and CNAG_05199 (green) predicted dimer. **E,** Key residues involved in NBD-NBD (1), NBD-SBD (2,3) and SBD-SBD (4) interactions. Hydrogen bonds are depicted in black. **F,** Structure prediction of Hsp90-CNAG_05199 dimer, with inset figures highlight key residues of Hsp90 (red) and CNAG_05199 (green) forming hydrogen bonds (black).

As Hsp70s frequently associate with other Hsp70 and Hsp90 proteins to perform chaperone activities, we further investigated the structural feasibility of the interactions of CNAG_05199 with both Hsp70 and Hsp90, as inferred from SEC-MS and STRING analyses. By applying AlphaFold 3.0 modelling to these protein complexes, we gained insight into the domains and regions involved in the predicted interactions. Starting with the CNAG_05199–Hsp70 complex, we observed high-confidence prediction scores (ipTM: 0.66, inter-chain PAE: 3.21–3.83 Å) with four distinct binding regions that formed large interfaces. In this complex, Hsp70 assumed a docked, ATP-bound conformation wrapping around a linear, ADP-bound CNAG_05199, with the interaction primarily mediated by NBD–SBD contacts **(Figures 8D and 8E)**. We then examined the CNAG_05199–Hsp90 complex, which was also confidently predicted (ipTM: 0.68, inter-chain PAE: 2.1–2.2 Å), but displayed a different binding configuration. In this complex, CNAG_05199 binds to the N-terminal and middle domains of *C. neoformans* Hsp90 via its NBD in the linear, undocked ADP-bound state, creating two large, buried interfaces **(Figure 8F).**

## Discussion

Our study unravels global molecular interaction networks of *C. neoformans* biology by applying an unbiased size fractionation approach coupled with high-resolution mass spectrometry. We aimed to define interactions between fungal proteins across the secretome and cellular proteome with roles in regulating cryptococcal virulence, with an emphasis on thermotolerance. As with other SEC-MS studies, our approach does not require *a priori* knowledge of PPIs within *C. neoformans* to explore new biological connections and reveals uncharacterized proteins with putative roles in heat shock regulation. Computational prediction analyses combined with experimental growth and thermotolerance assays confirm identification of heat shock associated proteins. Overall, we demonstrate the application of SEC coupled to mass spectrometry for providing new biological insights into the functional roles of prioritized targets within *C. neoformans*.

Our SEC findings demonstrate the complexity of the cryptococcal secretome, highlighting the presence of both extracellular and cell-associated proteins. Beyond potential contamination resulting from cell lysis, the detection of cell-associated proteins, such as 60S ribosomal protein L17 and ADP/ATP carrier, may reflect their export through extracellular vesicles (EVs), as demonstrated in previous studies (33,34). Given that cryptococcal EVs contain diverse protein cargo and can modulate host immune responses during fungal infection, their presence may represent an important mechanism for delivering intracellular proteins to the extracellular environment and influencing host–pathogen interactions (34–37). Moreover, among the secretome-associated proteins detected, HSPs and CipC have established roles in fungal virulence (34, 35). Notably, CipC modulates host responses in both *in vitro* and *in vivo* infection models, affecting host mortality and promoting fungal survival (37). Thus, the detection of intracellular and virulence-associated proteins expands the characterization of the cryptococcal secretome and supports a potential role for protein release in fungal adaptation and modulation of the host environment.

Despite SEC-MS providing unbiased and valuable insights into the composition and distribution of the cryptococcal secretome and proteome, our experiment also demonstrated its resolution limitation; while several proteins were concentrated within a limited number of fractions, others were distributed across multiple fractions. The complex nature of the samples (i.e., total cell extractions), may have contributed to overlapping elution profiles, causing fractionation bleed-through and peak widening that can hinder interpretation (39,40). Serial SEC could improve fractionation resolution, but at the cost of disrupting native protein complexes due to sample preparation protocols (41–43). Moreover, as SEC separates proteins based on their native states, fractions do not correspond directly to “size”; therefore, we can observe variable molecular weights within one fraction as well as a single protein being detected across fractions due to different molecular states (i.e., monomer vs. oligomer) (39–41,43). Additionally, proteins may be lost in the fractionation process due to non-specific adsorption, causing low-abundance molecules to fall below the mass spectrometry limit of detecton threshold and decrease comprehensiveness (40). Nevertheless, the use of analytical tools (e.g. STRING and AlphaFold) can assist with overcoming the constrained interpretation of protein distributions across fractions and unravel true interactors.

In this study, we applied STRING physical subnetwork to analyze SEC-MS proteome eluates, which revealed that large-subunit ribosomal and mitochondrial proteins were highly interconnected. Interestingly, despite their strong interplay during protein translation (44,45), physical interactions among mitochondrial and ribosomal proteins have not been widely described. Here, we predicted interactions between tricarboxylic acid cycle proteins and ATP synthases, supporting previous data observed in bacteria and mammals (46,47). Therefore, our results contribute to unravelling the molecular mechanisms involved in standard biological processes in *C. neoformans* that have not yet been widely reported for fungi.

In addition to identifying complexes in cryptococcal metabolism, our cellular proteome network profiling analyses also shed new light on virulence-associated factors, including thermotolerance. Notably, we predicted the interaction of the uncharacterized protein CNAG_00287 and Hsp70, a putative drug target in fungal infections that was recently associated with reducing antimicrobial resistance in *Mycobacterium tuberculosis* (7,48). Based on our sequence and structure analyses of CNAG_00287, we proposed its role as an SGT protein, a known enhancer of heat-shock ATPase activity associated with drug resistance (49,50), specifically through the conserved TPR carboxylate clamp – Hsp70 C-terminal mechanism (51). Interestingly, mutations in the interacting interface of the TPR-C-terminal in Hsp70 and Hsp90 increased the susceptibility of *A. fumigatus* to caspofungin, an antifungal that is inherently ineffective against *Aspergillus* spp. and *C. neoformans* (52,53). Thus, our findings support the categorization of CNAG_00287 as an SGT protein (SGT2) and highlight the importance of its interaction with Hsp70 as a promising candidate for drug target and repurposing.

Further examination of the role of CNAG_00287 in heat shock response revealed its biological importance in thermal resistance. We anticipated that deleting CNAG_00287 would disturb Hsp70 activity and therefore, affect thermotolerance; however, this phenotype was not confirmed under rich (i.e., YPD) *in vitro* growth conditions. Conversely, cryptococcal growth at 37 °C in YNB (i.e., nutrient-limited conditions) was impaired upon deletion of CNAG_00287. Since Hsp70 requires ATP for its activity, we also hypothesized that, provided cells had sufficient nutrients, other co-chaperones could compensate for the absence of CNAG_00287 (7); hence, no growth impact due to temperature changes was observed under rich conditions. However, nutrient deprivation weakened this compensation, leading to significant growth impairment at higher temperatures. A similar phenotype was also observed in *E. coli dnaK* deletion strains, in which starvation resulted in loss of thermotolerance (54). Moreover, insufficient nitrogen availability may decrease the cell’s ability to withstand stressful environments, including high temperatures (55,56), which could have accentuated the decrease in thermotolerance in CNAG_00287Δ. As a result, our experiments confirm the importance of CNAG_00287 in cellular heat acclimatation.

Sequence and structural analyses also corroborated the role of CNAG_05199 in the heat-stress response pathway. CNAG_05199 showed high sequence and structural conservation with the Hsp70 family, placing this protein as a putative Hsp70. Moreover, CNAG_05199 was confidently predicted to form heterodimers with *C. neoformans* Hsp70 and Hsp90. Notably, dimerization of fungal Hsp70s are required for their chaperone activity and resistance to thermal stress (57), supporting the occurrence of the predicted interactions in *C. neoformans.* In the CNAG_05199-Hsp90 complex, both structures were ADP-bound and undocked, with CNAG_05199 NBD binding to the correspondent NBD and SBD in Hsp90 (58,59). Similarly, crystal structures of *E. coli* and human Hsp70-Hsp90 complexes also showed this interface pattern, which is essential for client maturation and ATPase activity in Hsp90 (60,61). Given that Hsp70 and Hsp90 play a role in antifungal resistance (52,57,62), regulating its activity may provide a therapeutic potential by compromising fungal stress responses and improving susceptibility to antifungal agents. Altogether, these results provide evidence for the interplay of CNAG_05199 within the HSP families and advance our understanding of the cryptococcal stress response pathway. Further experimental validation is required to determine whether CNAG_05199 has biological importance for thermotolerance, as herein demonstrated for CNAG_00287.

## Conclusion

This study establishes a pipeline for investigating protein-protein complexes within the cryptococcal cellular and secreted proteomes, revealing both established and novel biological processes. By integrating SEC-MS with computational predictions, we elucidated the functions of the previously uncharacterized proteins CNAG_00287 and CNAG_05199 in the stress response pathway. Functional analysis confirmed the regulatory role of CNAG_00287 in thermotolerance, which demonstrates this approach as a proof-of-principle for the unbiased identification of novel virulence-associated proteins *in C. neoformans*. Ultimately, this work advances understanding of the cryptococcal interactome and underscores the potential of this method for investigating host-pathogen interactions.

## Author Contributions

M.dS. & J.G.-M. conceptualized and designed the study. M.dS., M.S. & F.R.-D. performed experiments, including protein extractions and sample preparations, proteomics sample preparation and mass spectrometry measurements. F.R.-D. & A.D. supported mass spectrometry experiments M.dS. & J.G.-M. analyzed data and generated figures. M.dS. & J.G.-M. wrote the first manuscript draft. All authors contributed to manuscript preparation and have read and approved the submitted manuscript.

## Acknowledgements

The authors would thank Laura Siedel from Dr. Cesar Khursigara lab at the University of Guelph for the technical support with the SEC. We also thank Dr. Mickael Leclerq for the support in the proteomics data analyses pipeline. The authors thank members of the Geddes-McAlister lab for helpful discussions and constructive comments on the study, especially Dr. Jason McAlister and Dr. Arjun Sukumaran.

## Data availability

The mass spectrometry data is available through the PRIDE Proteome Xchange Consortium: **Project accession:** PXD079582

Reviewer Token: kzsWRqP98pll

