## Supplemental file for "Mapping virulence-associated protein interaction networks reveals regulators of thermotolerance in *Cryptococcus neoformans*"

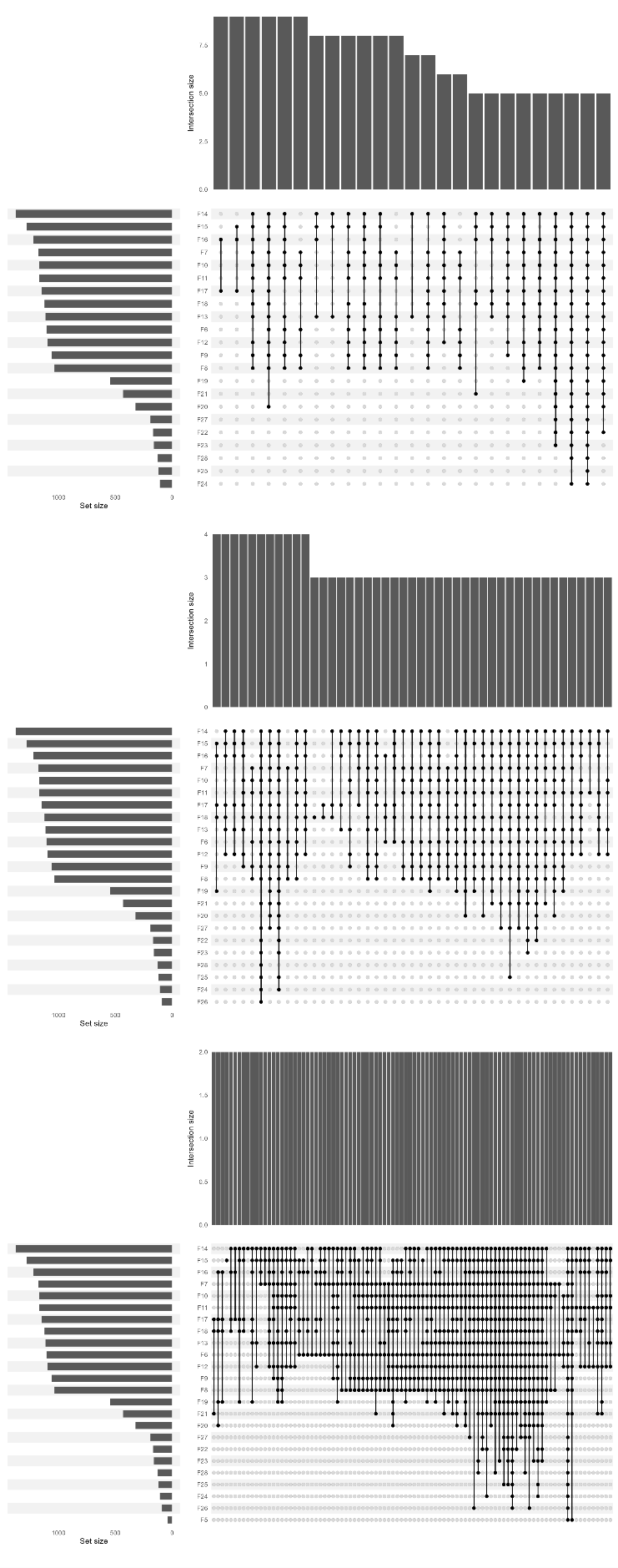

**Figure S1. Upset plot representing the number of proteins shared across *C. neoformans* proteome fractions.** Intersection size ranging from two to nine shared proteins. Experiment performed in biological quadruplicate.

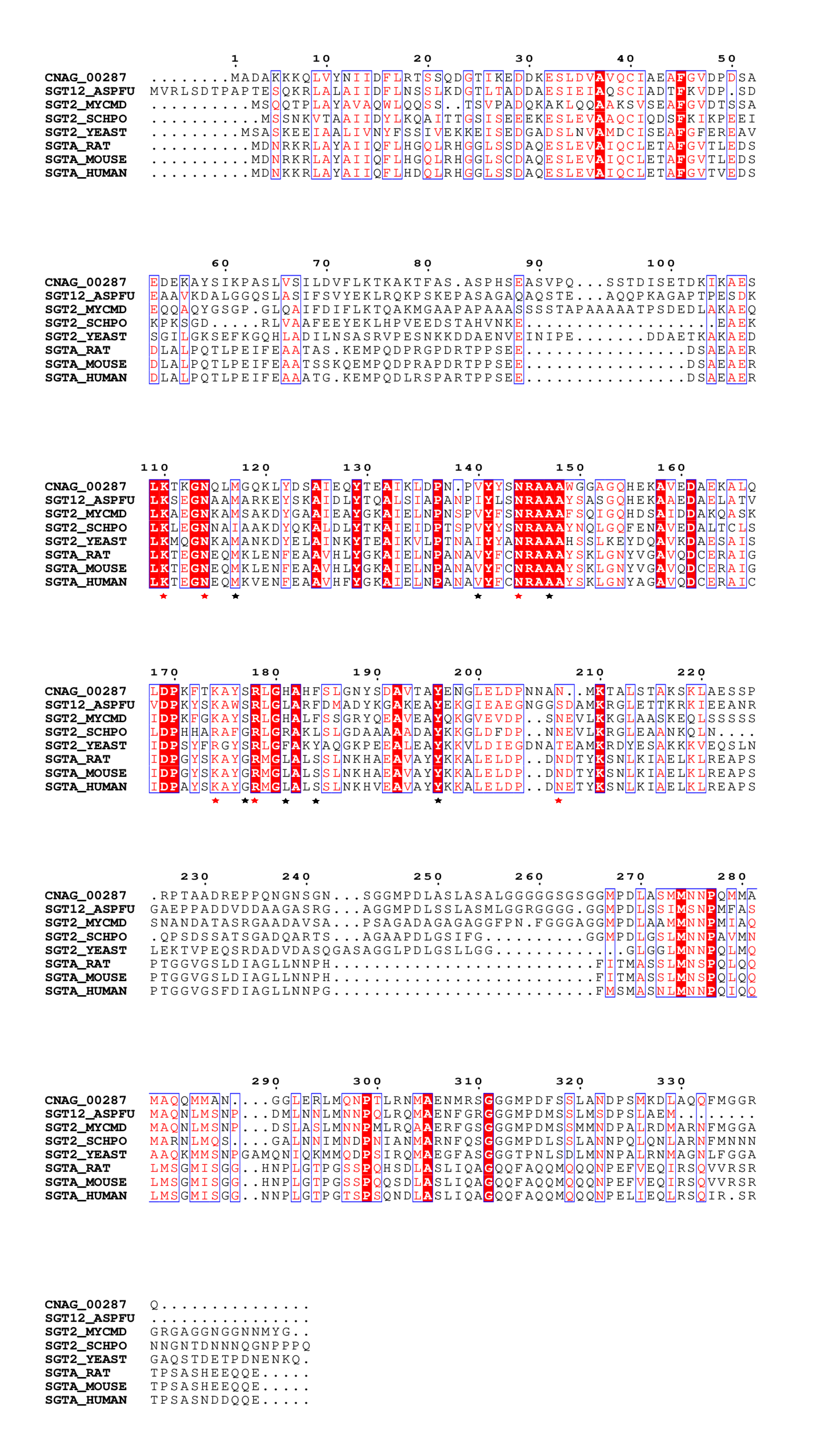

**Figure S2. Multiple sequence alignment of CNAG_00287 and closely related small glutamine-rich tetratricopeptide repeat-containing (SGT) proteins.** High conservation (red residues and background) in the TPR domain (stars). Red starts represent the carboxylate clamp, and black stars represent the residues responsible for stabilizing the interaction. SGTA_RAT: *Rattus norvegicus;* SGTA_MOUSE: *Mus musculus;* SGTA_HUMAN: *Homo sapiens;* SGT2_YEAST: *Saccharomyces cerevisiae;* SGT2_SCHPO: *Schizosaccharomyces pombe;* SGT12_ASPFU: *Aspergillus fumigatus;* SGT2_MYCMD: *Mycosarcoma maydis*.

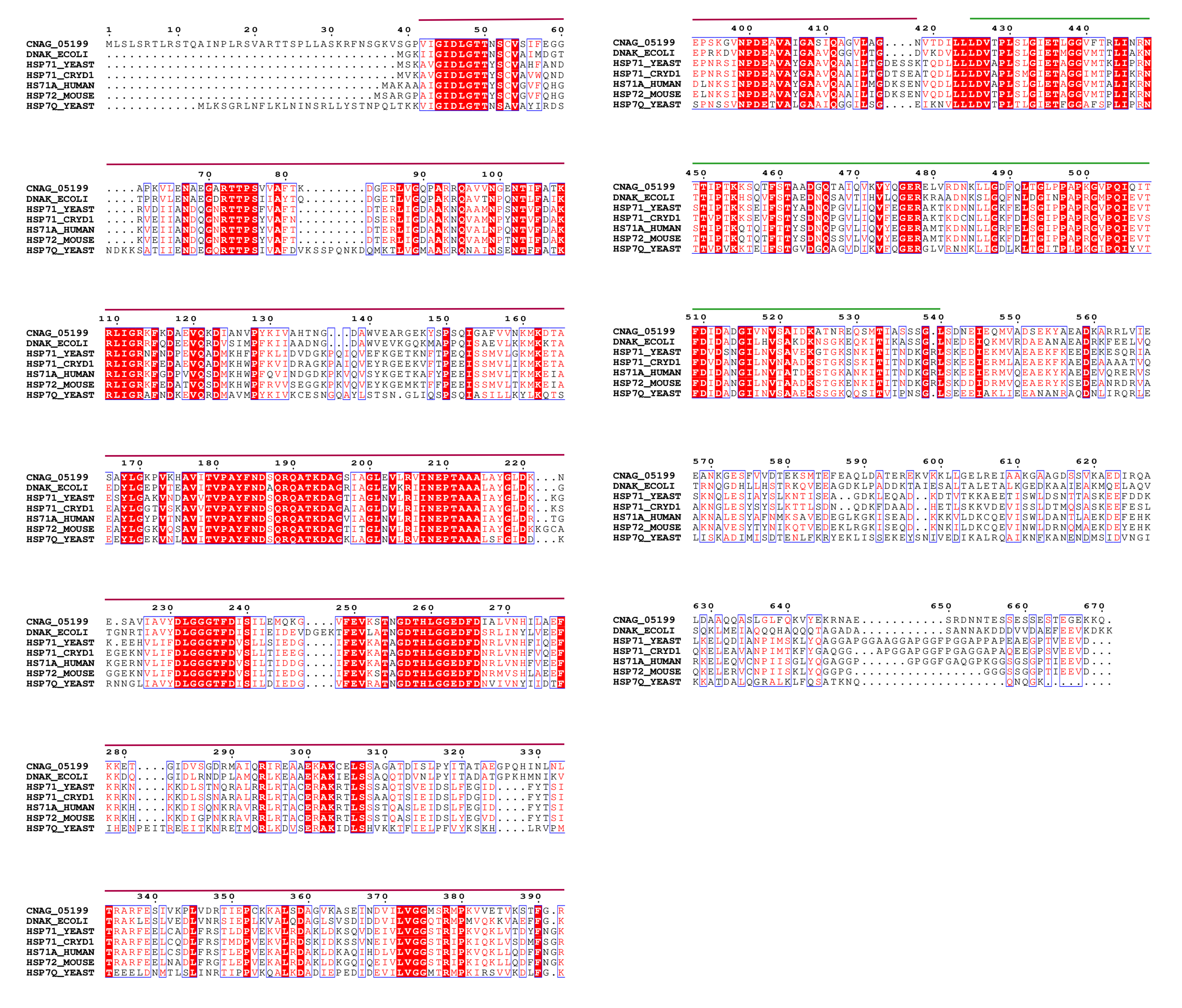
 **Figure S3. Multiple sequence alignment of CNAG_05199 and closely related Hsp70.** High conservation (red residues and background) in the nucleotide-binding domain (purple) and substrate-binding domain (green). DNAK_ECOLI: *Escherichia coli;* HSP7Q_YEAST and HSP71_YEAST: *Saccharomices cerevisiae,* mitochondrial and cytoplasmatic; HSP71_CRYD1: *Cryptococcus neoformans,* cytoplasmatic; HS71A_HUMAN: *Homo sapiens*; HSP72_MOUSE: *Mus musculus.*
